# CDK12/CDK13 inhibition disrupts a transcriptional program critical for glioblastoma survival

**DOI:** 10.1101/2023.07.14.548985

**Authors:** Silje Lier, Solveig Osnes Lund, Anuja Lipsa, Katrin B. M. Frauenknecht, Idun Dale Rein, Preeti Jain, Anna Ulrika Lång, Emma Helena Lång, Niklas Meyer, Aparajita Dutta, Santosh Anand, Gaute Johan Nesse, Rune Forstrøm Johansen, Arne Klungland, Johanne Egge Rinholm, Stig Ove Bøe, Ashish Anand, Steven Michael Pollard, Simone P. Niclou, Mads Lerdrup, Deo Prakash Pandey

## Abstract

Glioblastoma is the most prevalent and aggressive malignant tumor of the central nervous system. With a median overall survival of only one year, glioblastoma patients have a particularly poor prognosis, highlighting a clear need for novel therapeutic strategies to target this disease. Transcriptional cyclin-dependent kinases (tCDK), which phosphorylate key residues of RNA polymerase II (RNAPII) c-terminal domain (CTD), play a major role in sustaining aberrant transcriptional programs that are key to development and maintenance of cancer cells. Here, we show that either pharmacological inhibition or genetic ablation of the tCDKs, CDK12 and CDK13, markedly reduces both the proliferation and migratory capacity of glioma cells and patient-derived organoids. Using a xenograft mouse model, we demonstrate that CDK12/13 inhibition not only reduces glioma growth *in vivo*. Mechanistically, inhibition of CDK12/CDK13 leads to a genome-wide abrogation of RNAPII CTD phosphorylation, which in turn disrupts transcription and cell cycle progression in glioma cells. In summary, the results provide proof-of-concept for the potential of CDK12 and CDK13 as therapeutic targets for glioblastoma.

**Significance statement:** Glioblastoma is a common, aggressive, and invasive type of brain tumor that is usually fatal. The standard treatment for glioblastoma patients is surgical resection, radiotherapy, and chemotherapy with DNA-alkylating agents, and unfortunately current treatments only extend overall survival by a few months. It is therefore critical to identify and target additional biological processes in this disease. Here, we reveal that targeting a specific transcriptional addiction for glioma cells by inhibition of CDK12/CDK13 disrupts glioma-specific transcription and cell cycle progression and has potential to provide a new therapeutic strategy for glioblastoma.

## Introduction

With a median overall survival of 13.9 months and a 5-year survival rate of only 5.3%, IDH-wildtype Glioblastoma is the most prevalent and aggressive tumor of the central nervous system (1, 2). The standard of care for glioblastoma patients is surgical resection followed by radiotherapy and chemotherapy with temozolomide (TMZ), which only increases overall survival by a few months (3, 4). It is therefore critical to identify new specific vulnerabilities of glioblastomas that can be targeted pharmacologically.

Aberrant transcriptional programs are key to development and maintenance of cancer cells, and consequently cancer cells are often hypersensitive to the targeting of the transcriptional machinery (5). Glioblastoma propagation and resistance to existing therapies are driven by a subset of stem-like cells, which depend on neurodevelopmental transcription factors (TF) to maintain a specific transcriptional program and sustain proliferation (6, 7). As TFs can be functionally redundant or difficult to inhibit by small molecules, inhibition of the core transcriptional machinery offers an attractive alternative way to disrupt the transcriptional addictions of cancer cells.

RNAPII-dependent transcription is generally required for the transcriptional programs that sustain specific cell lineages and identities, including that of glioblastomas. The transcription cycle of RNAPII is regulated by a set of tCDKs, including CDK7-CDK13, that phosphorylate the RNAPII CTD and facilitate key steps of transcriptional initiation and elongation(8). CDK7 is involved in regulating transcriptional initiation by phosphorylating serine-5 (pSer5) of the RNAPII CTD (9). CDK9, CDK12, and CDK13 phosphorylate serine-2 (pSer2) of RNAPII CTD regulating transcriptional elongation (10, 11). tCDKs are attractive therapeutic targets (8), and several highly specific inhibitors were recently reported (12), including the CDK9-inhibitor NVP-2, the allosteric CDK7-inhibitor THZ1, and the allosteric CDK12/13-inhibitor THZ531. These inhibitors target a cysteine residue outside the kinase domain, thereby resulting in much higher specificity (11, 13). THZ1-mediated CDK7 inhibition leads to loss of RNAPII phosphorylation mainly at Ser5 and has anti-cancer properties in adult and pediatric glioma cells (14–16). THZ531 treatment reduces RNAPII phosphorylation mainly at Ser2, and can reduce neuroblastoma, osteosarcoma, and Ewing sarcoma proliferation (17, 18). Furthermore, the specific CDK12/CDK13 inhibitor SR-4835 reduces proliferation of triple negative breast cancer cells (19).

Here we explore whether tCDK inhibitors can inhibit glioma cell proliferation. Using THZ531 and SR-4835 that inhibit CDK12/CDK13 and RNAPII pSer2 ^11,19^, we demonstrate that glioma cell proliferation is specifically and strongly reduced due to loss of RNAPII phosphorylation, transcriptional shutdown, and disruption of a glioblastoma-specific transcriptional program. Finally, we demonstrate that the cell cycle progression and DNA replication are substantially affected in gliomas by CDK12/CDK13 inhibition. Altogether, our results illustrate that CDK12/CDK13 inhibition can provide a promising therapeutic alternative for the treatment of glioblastoma.

## Results

### Inhibition of CDK12/CDK13 arrests glioblastoma cell proliferation and migration

To identify tCDKs affecting glioblastomas proliferation, we first performed survival analysis using GlioVis (20), revealing that glioblastoma patients with higher CDK13 expression have significantly poorer overall survival (HR = 0.66; p = 0.0058). In contrast, no significant correlation was observed for other tCDKs (Supplementary Figure 1A), although all tCDKs were expressed in GSCs (Supplementary Figure 1B). We therefore examined the effect of selected tCDK inhibitors on the survival of a panel of glioma-patient derived stems cells (GSCs) and control cells using dose response analyses (Supplementary Figure 1, D-F). We found GSCs to be particularly vulnerable to two small molecule CDK12/CDK13 inhibitors THZ531 and SR-4835 (11, 19) (Figure 1, A and B, Supplementary Figure 1, F and G). Furthermore, the effect was independent on the presence of serum in the cell culture media (Supplementary Figure 1C). Importantly, all GSCs tested were sensitive to THZ531 and SR-4835 treatment, and their IC_50_ values ranging from 20 to 200 nM were substantially lower than those of cells from other cancer sub-types (Figure 1B, Supplementary Figure 1F). In agreement with a recent study reporting that THZ531 inhibited proliferation of liver cancer cells (21), human hepatoma HepG2 cells were also sensitive to THZ531. Moreover, 100 nM and 500 nM THZ531 led to a strong reduction in proliferation and colony formation for the GSCs, but not for Hela cells (Figure 1C and D, Supplementary Figure 1G). To further validate the results obtained using inhibitors, we investigated the effect of genetic ablation of CDK12/CDK13 on the proliferation of GSCs using a CRISPR/Cas9-based competition assay. Positive control single guide RNAs (sgRNA) against MCM2, RPS19, and CDK9 inhibited glioma cells, whereas a negative control had no effect. Importantly, each of three independent sgRNAs targeting CDK12, or CDK13, revealed that genetic ablation of these targets significantly inhibited the glioma cell proliferation (Figure 1E).

**Figure 1.**
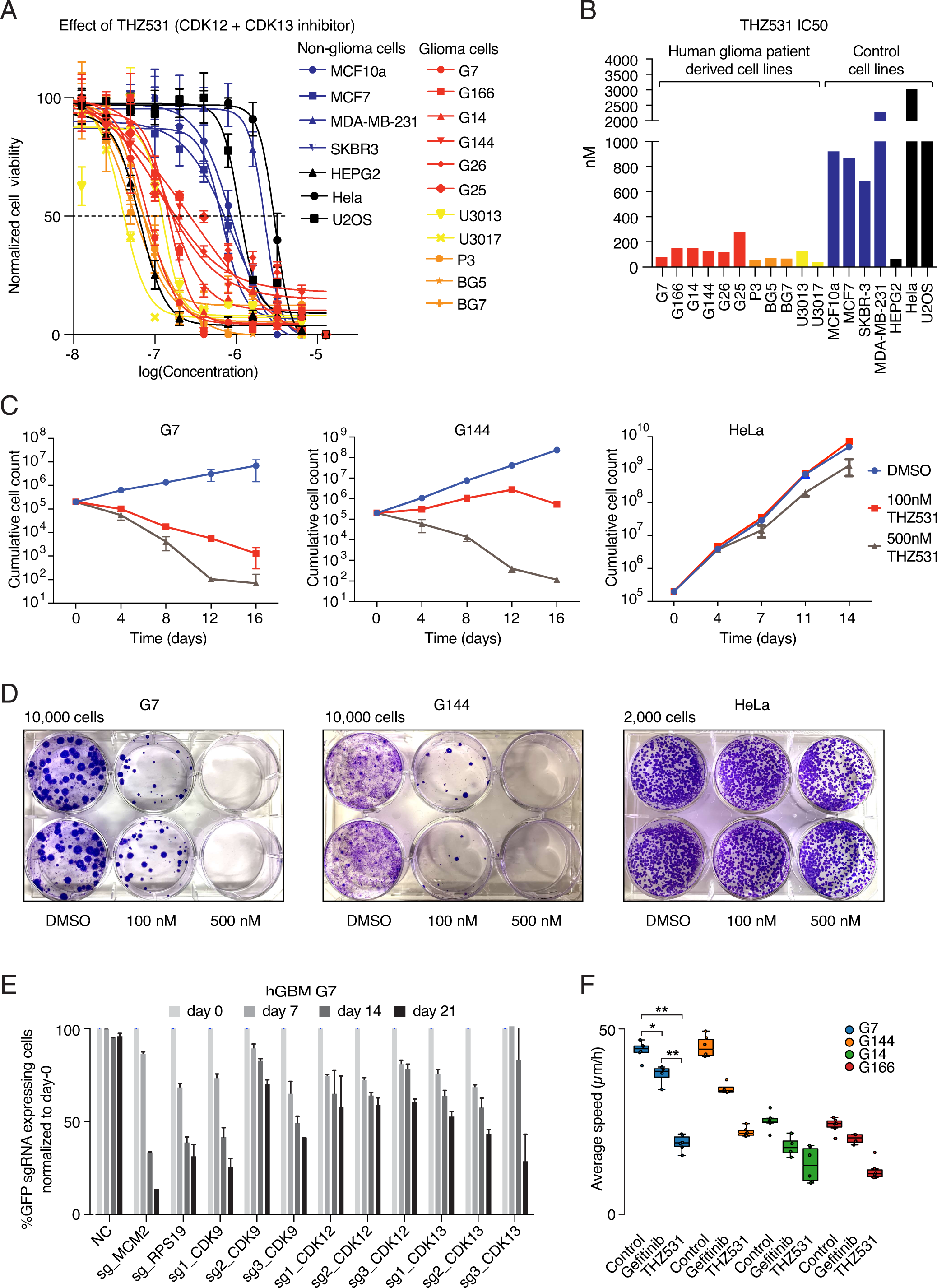
Inhibition of CDK12 and CDK13 specifically affects proliferation of glioma cells. (**A**) Dose-response curves from MTT assays eleven high-grade glioma and seven non-glioma cell lines treated with THZ531. Data represent mean ± SD of three replicates. (**B**) Bar graph showing IC50 values from the MTT assays in (A). (**C**) in vitro cell proliferation assay of the GSCs G7 and G144, and HeLa cells treated as indicated. Data represent mean ± SD of three replicates. (**D**) Clonogenic survival assay of G7, G144 and HeLa cells treated as indicated. (**E**) Competition assays of sgRNAs targeting CDK9, CDK12 and CDK13 as well as positive controls (essential genes, MCM2 and RPS19). Non-targeting sgRNA (NC) was used as negative control. (**F**) Boxplot representing average migration speed of four high-grade glioma cells treated as indicated. The EGFR inhibitor Gefitinib was used as positive control. Each data-point boxplots represents average migration speeds in 4-6 acquired time-lapse movies. The boxplot is representative of three independent experiments. *p < 0.01, **p < 0.001.

Cell migration is central for the invasive capacity of malignant gliomas (22), and we next assessed the effect of THZ531 on GSC migration compared to the positive control Gefitinib (23). High-content live-cell microscopy revealed that THZ531 significantly reduced migration of G7, G144, G14 and G166 GSCs and even exceeded the ability of Gefitinib to inhibit GSC migration (Figure 1F, Supplementary Figure 1H, Supplementary movies 1 and 2). These findings demonstrate that THZ531 strongly inhibits proliferation and migration of glioma cells.

### CDK12 is expressed in human glioblastoma tissue and inhibition of CDK12/CDK13 compromises *ex vivo* glioblastoma proliferation

Immunohistochemistry for CDK12 was performed on glioblastoma tissue (n = 5 patients) and on CNS tissues without glioblastoma (n = 2 patients), see Supplemental Table 1 for details on patient material. In control CNS tissue, no unequivocal nuclear CDK12 expression was detected, while a heterogeneous distribution of nuclear CDK12 expression was visible in glioblastoma tissue, ranging from weak, to moderate, to strong expression (representative pictures are provided in Figure 2A, bottom panel).

**Figure 2.**
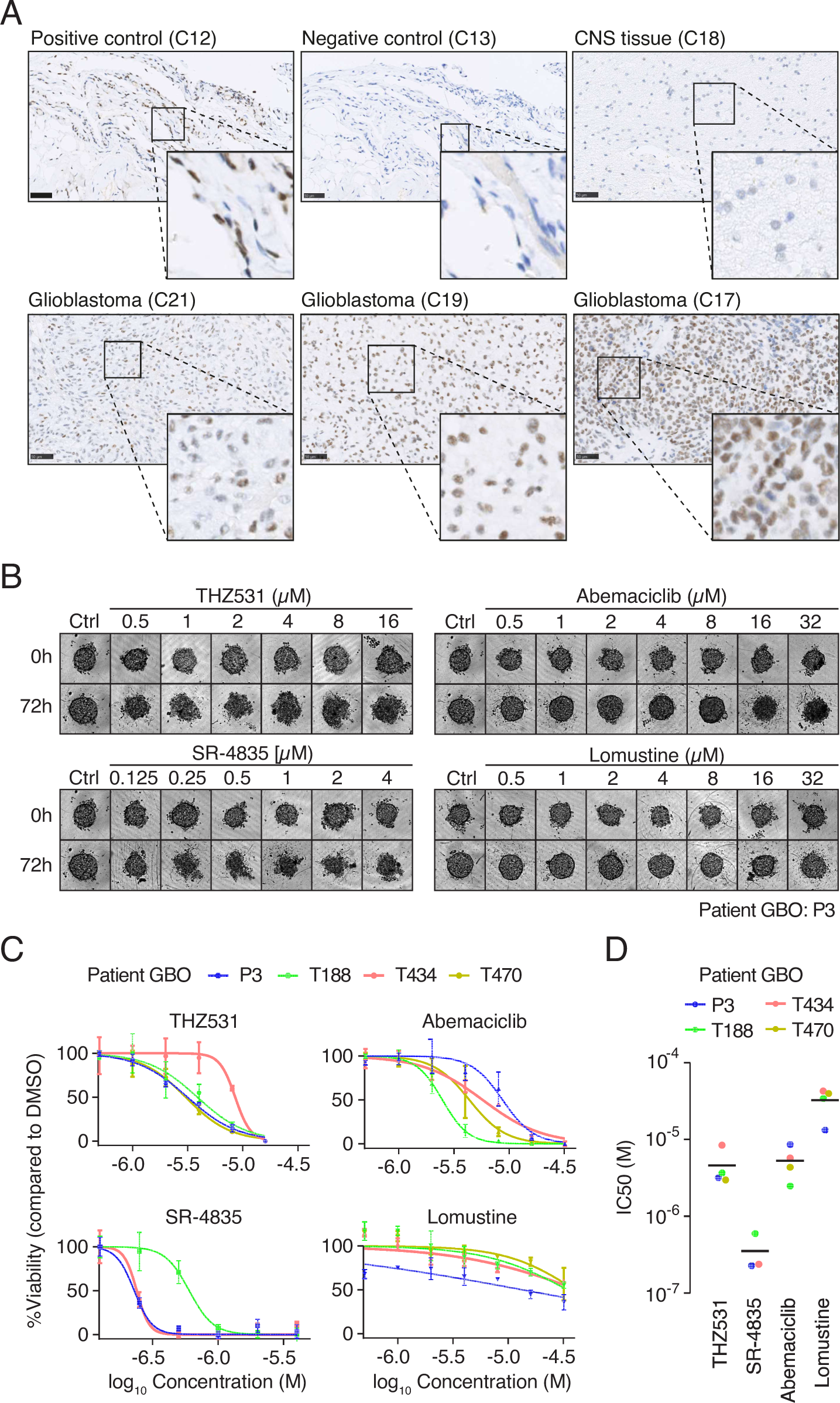
CDK12 is expressed in resected glioblastoma tissue of human patients and inhibition of CDK12/CDK13 compromises ex vivo glioblastoma proliferation. (**A**) Representative images of CDK12 immunohistochemistry in control and glioblastoma tissue. The top row shows control stainings whereas the bottom row left to right reflect CDK12 expression in three different glioblastoma patients (for information on patients, see Supplementary Table 1). Scale bar 50 μM. (**B**) Representative image showing effect of different inhibitors with indicated doses on glioblastoma organoids. (**C**) Dose-response curve showing data from glioblastoma organoids treated with inhibitors as indicated and measured for cell viability 72 h post treatment. (**D**) Dot-plot showing IC50 values from the assays in (**C**).

We next assessed the effect of CDK12/CDK13 inhibitors on ex vivo 3D glioblastoma patient-derived organoids (GBOs) that were reformed from isolated single tumor cells as previously described (24, 25). Four glioblastoma patient-derived tumor cells were selected, exhibiting a typical range of key glioblastoma genetic alterations including deletion of CDKN2A/B, amplification of CDK4/6 and EGFR, mutations of TP53, PTEN, PIK3CA and EGFR (Supplementary Figure 2B). Furthermore, in addition to the CDK12/CDK13 inhibitors, THZ531 and SR-4835, we included Abemaciclib, a CDK4/6 inhibitor and Lomustine, both inhibitors of clinical significations for glioblastoma treatment to benchmark the effect of CDK12/CDK13 inhibition. THZ531 and SR-4835 treatment strongly affected the morphology and structure of GBOs (Figure 2B) and inhibited the proliferation of GBOs (Figure 2, C and D). The effect of THZ531 treatment on inhibition of GBO was comparable to Abemaciclib and Lomustine where SR-4835 was a more potent inhibitor of GBO proliferation compared to Abemaciclib and Lomustine (Figure 2, C and D), a consistent finding observed in GSCs (Supplementary Figure 2, C and D).

### Inhibition of CDK12/CDK13 reduces *in vivo* tumor growth

To investigate the effect of CDK12/CDK13 inhibitors can reduced tumor burden in vivo, we used CDK12/13 inhibitor SR-4835 because it has been tested against triple-negative breast tumor xenografts in mice (19), while THZ531 has not yet been used for *in vivo*. We determined that in mice, the maximum tolerated dose of SR-4835 was 20 mg/kg. In addition, we discovered that SR-4835 was undetectable in mouse brains when sampled 24 h post-dose, suggesting that it cannot cross the blood-brain barrier (BBB). Therefore, we used a mouse subcutaneous xenograft model based on U87-MG cells to test the efficacy of the CDK12/13 inhibitor SR-4835, and compared it to TMZ treatment. SR-4835 reduced *in vitro* U87-MG proliferation with similar IC50 as we have observed for GSCs (Figure 3, A and B). Nine days after injection with U87-MG cells, mice were dosed with SR-4835 or TMZ for two weeks (Figure 3C). Growth of subcutaneous tumors was strongly inhibited by 20 mg/kg SR-4835, 5 mg/kg or 2 mg/kg TMZ (Figure 3C). Importantly, the constant body weight of the mice indicated that all treatments were well tolerated. Finally, we tested the effect of SR-4835 on a large panel of cancer cell lines encompassing pancreatic, ovarian, uterine and prostate cancer, and found that GSCs were most sensitive to CDK12/CDK13 inhibition (Figure 3D). Altogether, we found that inhibition of CDK12/13 has a strong and specific effect on glioma growth, which compares favorably with the existing treatment.

**Figure 3.**
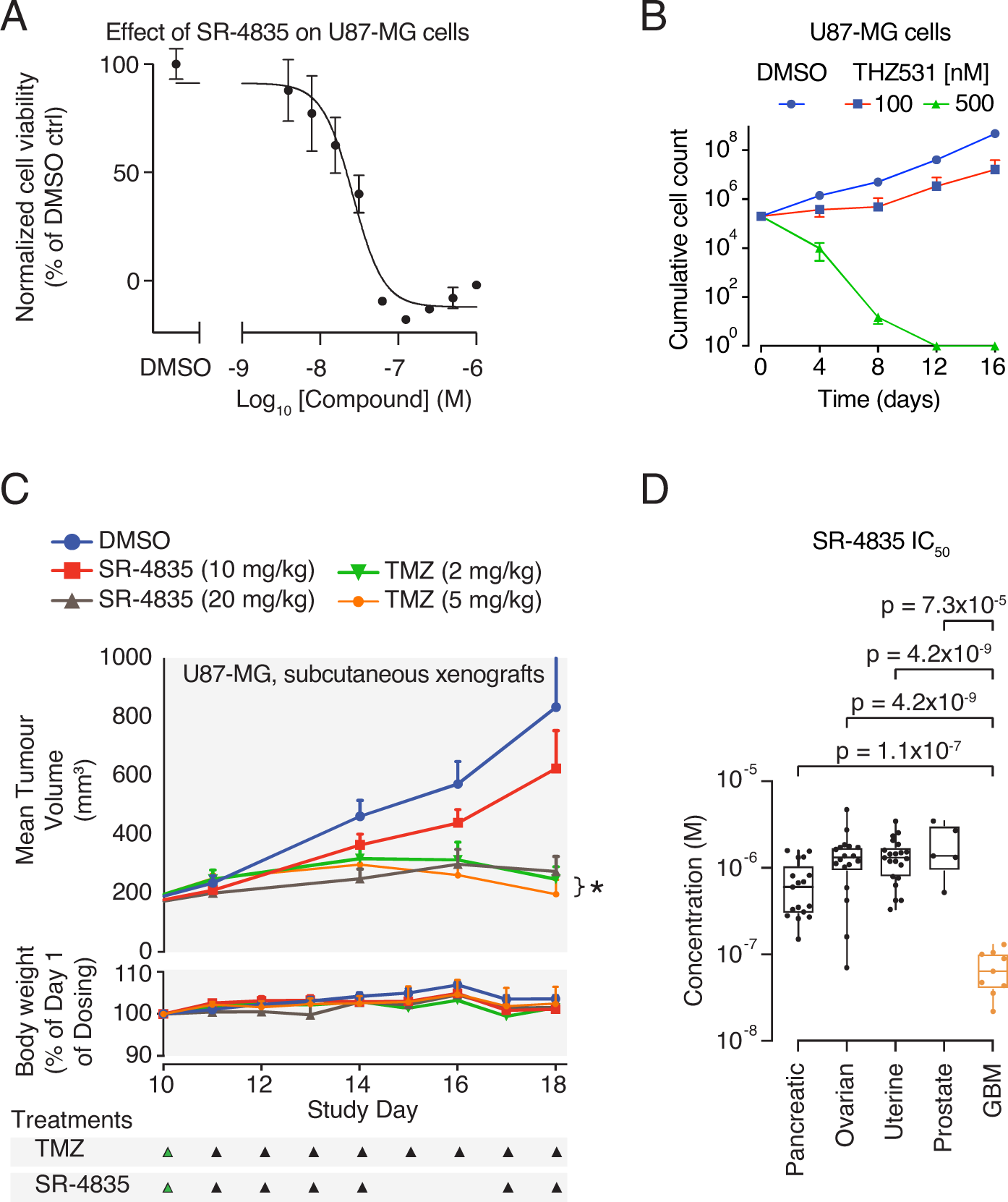
Inhibition of CDK12/CDK13 reduces tumor burden in vivo. (**A**) Dose-response curve for U87-MG glioma cells treated with SR-4835. Data represent mean ± SD of three replicates. (**B**) in vitro cell proliferation assay U87-MG cells treated as indicated. Data represent mean ± SD of three replicates. (**C**) Mean tumor volume and relative body weight of mice over time for the indicated treatments with the dosage shown in the bottom. Significance of the combination treatment is indicated with an asterisk. (**D**) Box-plot showing IC50 values for dose response of SR-4835 inhibition on a panel of cancer cell lines. p-values are from two-sided t-tests Benjamini-Hochberg corrected for multiple testing.

### Inhibition of CDK12/CDK13 leads to global loss of RNAPII CTD phosphorylation and nascent mRNA synthesis in GSCs

We next analyzed the phosphorylation level of key residues in the RNAPII CTD following THZ531 treatment. Using 500 nM THZ531 as in previous studies of other cell types (11, 18, 26, 27), we found that 6 h of treatment almost completely abolished Ser2 phosphorylation and strongly affected pThr4 in GSCs G7 cells, while the total RNAPII levels remained unchanged up to 24 h (Figure 4A). In contrast, no substantial changes were observed in the levels of phosphorylated species of RNAPII within 48 h of treatment for HeLa cells (Figure 4A).

**Figure 4.**
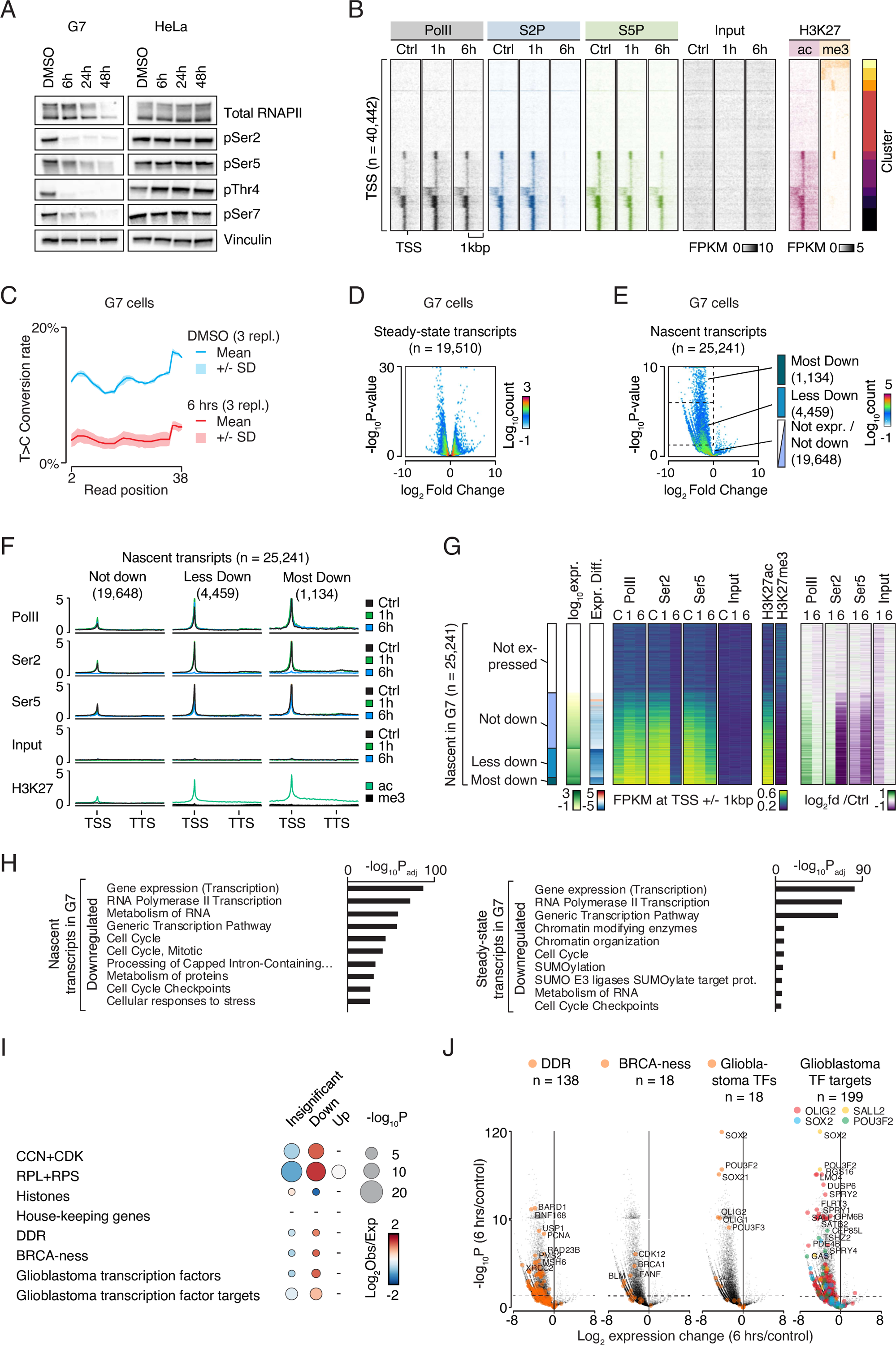
CDK12/CDK13 inhibition profoundly affects RNAPII phosphorylation and transcription in glioma cells. (**A**) Immuno-blot analyses from G7 and HeLa cells that were treated with vehicle, 500 nM THZ531 for 6 h, 24 h and 48 h for the various RNAPII CTD species. (**B**) Heatmaps of Cut&Run signal from RNAPII, RNAPII phosphorylation states, and histone modifications at k-means clustered unique TSSes +/-1 kbp. G7 cells were treated with DMSO and THZ531 for 1 h and 6 h. (**C**) Diagram showing T to C conversion in three replicates of SLAM-seq data from G7 cells treated with THZ531 or DMSO for 6 h. (**D**)-(**E**) ‘Volcano plots’ showing the overall transcriptional differences in G7 cells treated with THZ531 for 6h compared to DMSO control for steady-state (**D**) and nascent transcripts (**E**). X-axis shows the log2 fold differences. Y-axis shows the −log10 Benjamini-Hochberg corrected p-values. Dashed rectangles: populations of interest. (**F**) Graphs of average Cut&Run signal from RNAPII, RNAPII phosphorylation states, and histone modifications at and around TSS and TTS of genes. Levels are FPKM normalized. (**G**) Heatmaps illustrating transcription in relationship to Cut&Run data of RNAPII, RNAPII phosphorylation states, and histone modifications at TSSes. Groups from Figure 4E (**H**) Bar diagrams of Benjamini-Hochberg corrected −log10 p-values from most enriched gene ontology terms in nascent or steady state transcripts down-regulated after 6h of THZ531-treatment in G7 cells. (**I**) Bubble diagram of enrichment of gene groups within up-regulated, down-regulated or unchanged categories. P-values are Benjamini-Hochberg corrected). (**J**) Volcano plots as in Figure 4E with the indicated gene populations colored.

To investigate the effect of CDK12/CDK13 inhibition on genome-wide chromatin occupancy of total RNAPII, pSer2 and pSer5 in glioma cells, we employed the recently developed CUT&RUN technology (28). Total RNAPII occupancy transiently increased at genes with preexisting RNAPII at transcription start sites (TSSes) after 1 h of THZ531 treatment and were similar to those of control treated cells after 6 h of treatment (Figure 4B). pSer5 levels were considerably reduced post 6 h treatment, whereas genome-wide chromatin occupancy of pSer2 was strongly reduced following 6 h of treatment. Control markers H3K27ac and H3K27me3 were enriched at TSSes of actively transcribed and silent genes, respectively (Supplementary Figure 3A). In summary, these data show that THZ531 treatment strongly reduced genome-wide levels of phosphorylated RNAPII species.

To examine nascent and steady state transcription in THZ531-treated cells, we used SLAM-seq (29). In brief, uridines of nascent transcripts were labeled using a short pulse of 4-thiouridine (4sU) and isolated RNA was then exposed to iodoacetate, which converts incorporated uridine to cytosine. T->C conversion was used to identify nascent transcripts and compare the patterns of nascent and steady state transcription in THZ531-treated and control G7 cells. This established that THZ531 strongly suppressed nascent mRNA synthesis (Figure 4C). Indeed, exposure to THZ531 for 6 h caused a near total loss of newly-transcribed mRNAs and also had a strong impact on the composition of the steady state transcriptome, with thousands of mRNAs being significantly up- and down-regulated (Figure 4, D and E).

As expected, the expression of mRNAs showed a very high level of concordance with total RNAPII occupancy as well as RNAPII phosphorylation (Figure 4, F and G). In line with the observation that THZ531 nearly completely blocked nascent transcription, the most highly expressed transcripts with the highest levels of TSS-associated RNAPII, were also most strongly down-regulated by THZ531 (Figure 4, F and G). Gene Ontology analysis revealed profound consequences of THZ531-mediated inhibition of CDK12/CDK13, and the most strongly enriched functional categories of down-regulated nascent and steady state transcripts in THZ531-treated GSCs, G7 and G144, included transcription, cell cycle, and RNA metabolism (Figure 4H, Supplementary Figure 3C). Up-regulated steady-state transcripts were enriched in biological processes, such as translation and metabolism (Supplementary Figure 3C) in accordance with previous observations (17, 18). We also investigated selected sets of transcripts, including ribosomal, histones, cell cycle, house-keeping, DNA damage response (DDR), BRCA-ness factors, which are known to be dependent on CDK12 for their expression (18, 26), as well as the TFs that sustain the proliferation of glioma cells and their targets (Figure 4I). Nascent transcripts encoding cell cycle, DDR, BRCA or core glioma TFs and their target genes were significantly enriched among the down-regulated genes (Figure 4, I-J, Supplementary Figure 3D and 4A). In contrast, we found that nascent transcripts expressed from housekeeping genes were not significantly enriched or depleted among up-regulated, down-regulated, or unchanged genes.

Transcripts encoding key neurodevelopmental TFs, including OLIG2, POU3F2 and SOX2 are required for proliferation of glioma cells, were among the most strongly down-regulated transcripts (Figure 4J). The down-regulation or ablation of these TFs has been shown to strongly suppress proliferation of glioma cells (6). We therefore investigated change in their expression and occupancy of total, pSer2, and pSer5 forms of RNAPII following THZ531 treatment. All three genes were highly expressed in G7 cells and were marked by the presence of super-enhancer sites containing broad H3K27ac domains, as reported earlier (6, 7) (Supplementary Figure 5A). While pSer2 was nearly abolished at these genes and pSer5 strongly reduced, total RNAPII was also reduced following 6 h of CDK12/CDK13 inhibition. We observed a strong down-regulation in the nascent transcripts for these genes, which was validated using RT-qPCR. We found that the expression of key DDR genes was strongly down-regulated following THZ531 treatment as reported earlier (Supplementary Figure 5B). In addition, the expression of many glioblastoma-specific neurodevelopmental TFs including OLIG2, POU3F2 and SOX2 target genes was profoundly down-regulated (Supplementary Figure 5C), with Olig2 target genes being most strongly affected whereas expression of many, but not all house-keeping genes remained unaffected (Supplementary Figure 5D). In concert, these results demonstrate that THZ531-mediated inhibition of CDK12/CDK13 strongly suppresses expression of TFs (and their target genes) required for the glioblastoma transcriptional program.

### THZ531 treatment disrupts GSC cell cycle progression

G7 and G144 GSCs responded similarly to THZ531 and overlapping sets of differentially expressed transcripts were readily identified (Supplementary Figure 6A). Several functional categories related to the cell cycle were enriched among the most down-regulated nascent transcripts in THZ531-treated G7 cells and in the steady-state transcripts down-regulated in both G7 and G144 cells (Figure 4H, Supplementary Figure 3C). Publicly available data on cell cycle dependence of gene expression in HeLa and U2OS cells (30) were used to assess cell cycle-dependence of transcriptional changes following THZ531 treatment in GSCs. The results showed the strongest down-regulation of nascent transcripts in THZ531-treated G7 cells after S-phase (Supplementary Figure 6, B and C). Interestingly, prior to S phase, the steady state transcripts were both up- and down-regulated whereas after S-phase, the majority of the steady state transcripts were down-regulated (Supplementary Figure 6, B and C).

To further explore the effect of CDK12/CDK13 inhibition on the cell cycle of glioma cells, we used EdU incorporation to mark the cells in S-phase. Following 6 h THZ531 treatment, there was no EdU incorporation in both the G7 and G144 cells, indicating a lack of DNA synthesis (Figure 5, A and B) and this effect was stable up to 24 h. At the same time, we did not observe any change in EdU incorporation in Hela cells following THZ531 treatment (Supplementary Figure 6D). Furthermore, we noticed an increase of cells in both G1- and G2-phases following the pronounced loss of cells in active S-phase. We arrested cells in mitosis using nocodazole in combination with THZ531 treatment and found that GSCs were not entering mitosis, indicating blocked cell cycle progression (Figure 5, C and D). Furthermore, neither the frequency of apoptosis nor the amount of DNA damage changed after 6 h exposure to THZ531. However, after exposure to THZ531 for 24 h, apoptotic cells, DNA damage, γ-H2AX and cleaved PARP did increase (Figure 5, E-G). Expression of key cyclin genes, which are required for different phases of cell cycle was found to be severely down-regulated following THZ531 treatment (Figure 5H), explaining the arrested cell cycle progression. These data indicate that THZ531 impacts DNA synthesis and cell cycle progression of glioma cells, in addition to blocking nascent transcription and phosphorylation of RNAPII CTD.

**Figure 5.**
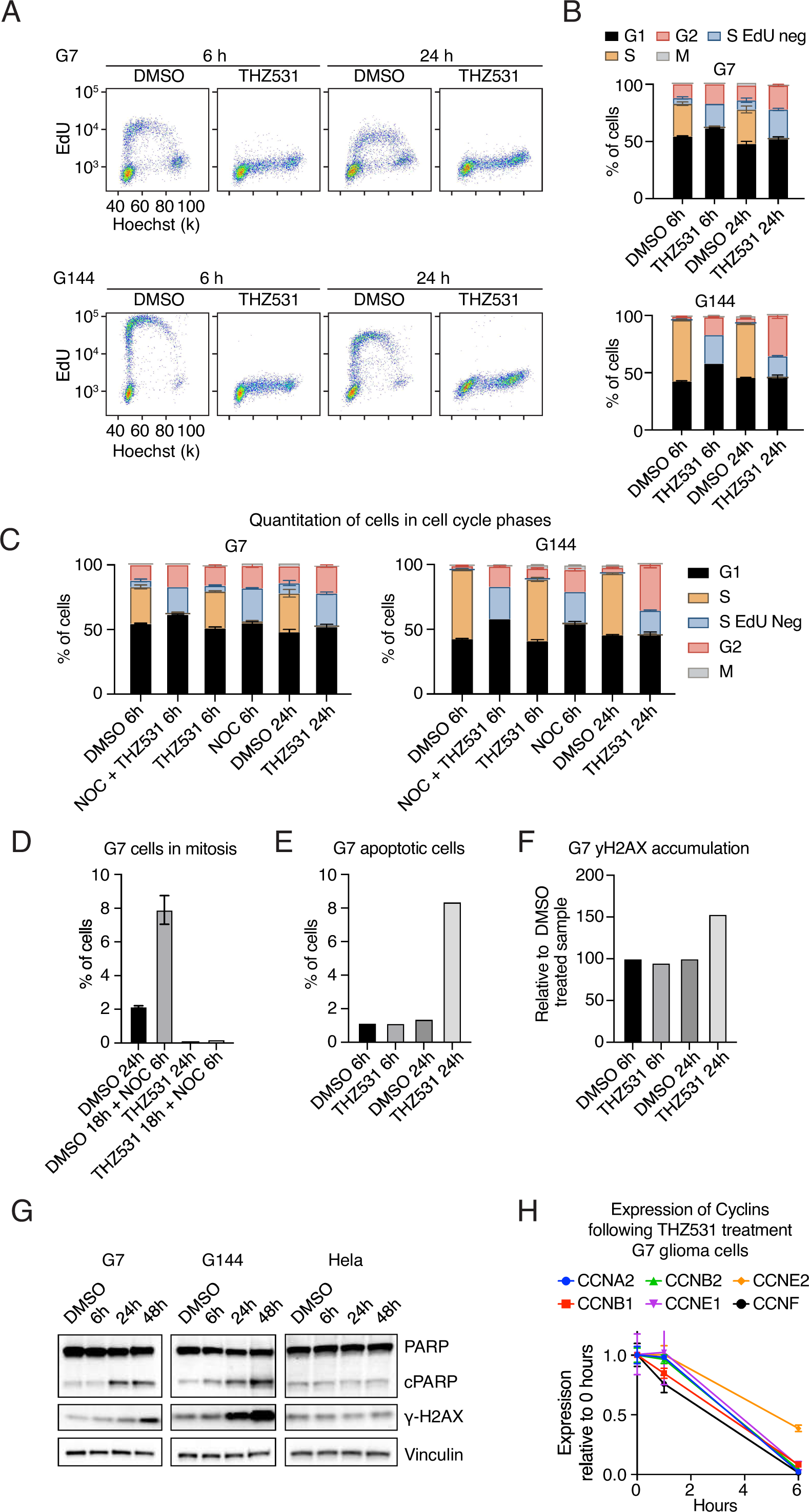
CDK12/CDK13 inhibition disrupts glioma cell cycle. (**A**) Cell cycle analysis in G7 and G144 cells after 6 and 24 h treatment with DMSO or 500 nM THZ531. 10 µM EdU was added 1 h prior to harvest. Dot plots show intensity of EdU relative to DNA content (Hoechst) in interphase cells (one representative replicate shown). (**B, C**) Bar diagrams of cell cycle distributions of G7 and G144 cells treated as indicated. Doses used: THZ531 (500 nM) Nocodazole (1 µg/ml). (**D**) Flow cytometry analysis of % of G7 cells in mitosis following 500 nM THZ531 ± 1 µg/ml Nocodazole treatment for 24h. Data represent mean ± SD of two replicates for (**B**)-(**D**). (**E**, **F**) Flow cytometric assays of apoptotic (**E**) of γH2AX accumulation (**F**) cells in G7 cells treated with 500 nM THZ531 for 6 h and 24 h. (**G**) Western blots of PARP and γH2AX levels in G7, G144 and HeLa cells treated with 500 nM THZ531. (**H**) RT-qPCR analysis of cyclin mRNA levels in G7 cells treated with 500 nM THZ531 for 1 h and 6 h. Data represent mean ± SD of two replicates.

## Discussion

In the present study, we identify an efficient and specific way to target and disrupt the transcriptional program required for glioma cell proliferation and migration. Specifically, we show how small molecule inhibitors targeting CDK12/CDK13, which phosphorylate RNAPII, strongly perturb the transcriptional landscape in GSCs. This leads to down-regulation of multiple glioblastoma-associated transcription factors and their targets, and subsequently a strong inhibition of GSC-proliferation, stalled cell cycle progression, and induction of cell death. The clear dependence of glioma cells on CDK12 and CDK13 for proliferation was validated using CRISPR/CAS9-mediated genetic ablation of CDK12 and CDK13. We also confirmed that CDK12 is well-expressed in human glioblastoma tissue ranging from weak to moderate to strong expression. Moreover, the CDK12/CDK13 inhibitors affected the morphology and reduced the survival of ex vivo GBOs, in a favorable manner compared to CDK4/6 inhibition and Lomustine, both of which are used to treat glioblastoma patients. While our results show that the CDK12/CDK13 inhibitor SR-4835 is well-tolerated in mice, it fails to cross the BBB. However, the observed reduced tumor burden in a subcutaneous xenograft mouse model, encourages further tests in more clinically relevant *in vivo* models as well as identification of novel CDK12/13 inhibitors that can pass the BBB.

The primary activity of CDK12 and CDK13 is to phosphorylate Ser2 of RNAPII CTD (8) and exposure to THZ531 rapidly and nearly completely blocks phosphorylation of RNAPII Ser2 and strongly reduces abundance of other RNAPII phospho-species in GSCs, with no similar effect in control cells. Furthermore, pSer2- and pSer5-RNAPII chromatin occupancy is strongly reduced in THZ531-treated glioma cells, while total RNAPII chromatin occupancy is not. In addition, exposure of glioma cells to THZ531 nearly abolishes nascent mRNA synthesis and causes large-scale changes in steady state mRNA expression. In particular, THZ531 strongly down-regulates expression of TFs and transcriptional programs essential for glioma cell proliferation (6).

We find that CDK12/CDK13 inhibition has a profound impact on expression of genes involved in transcription and cell cycle regulation. Concordantly, THZ531 rapidly suppresses active DNA replication and increases the proportion of cells in G1 and G2 in GSCs. However, apoptosis and DNA damage are observed after 24 h exposure to THZ531, but not after 6 h. These observations suggest that THZ531 compromises DNA synthesis, and that this subsequently may lead to DNA damage and apoptosis. At the same time, we find that the expression of key cyclin genes together with core DDR genes is profoundly down-regulated within 6 h treatment of THZ531, explaining the remarkable arrest of cell cycle progression. Glioblastomas have constitutive activation of the DDR pathway and show high genomic instability (31–33), and several key regulators of the DDR pathways, including BRCA1, are required for glioblastoma survival ^41^. We find that key DDR components are suppressed rapidly and strongly by CDK12/CD13 inhibition, providing a potential explanation for the remarkable sensitivity of GSCs compared to other cancer cells (Figure 3F).

Since the initial identification of THZ531 in 2016, there has been great interest in understanding the requirement and roles of CDK12/CDK13 in cancer cells. Furthermore, loss or mutation of CDK12 is reported for several cancers, including ovarian, breast and prostate cancers (34–36). These studies correlate loss of CDK12 with altered expression of core DNA damage response genes. Recent studies associated expression changes after loss of CDK12/CDK13 with gene length, expression level, and intronic polyadenylation cleavage of affected mRNAs (17, 18, 26). In agreement with other studies focusing on breast cancer and neuroblastomas (17–19, 26), we find that the inhibition of CDK12/CDK13 strongly down-regulated the expression of core DDR genes. Furthermore, our observations are in agreement with recent findings in K562 chronic myeloid leukemia cells, where inhibition of CDK12/13 result in a global loss of RNAPII CTD phosphorylation and extensive genome-wide transcriptional changes (27). Moreover, similar results are obtained when CDK12 is inhibited in HEK-293 cells (37). At higher concentrations, we observe that the proliferation of control cells is also affected by both CDK12/CDK13 inhibitors, THZ531 and SR-4835. Therefore, other cancer cells potentially require a higher dose of CDK12/CDK13 inhibitors to observe the global changes in RNAPII phosphorylation and subsequent shutdown of nascent transcription. It will be relevant for future work to understand the factors that govern the differences leading to sensitivity of CDK12/13 inhibition among different cancers.

Taken together, we report that inhibition of CDK12/CDK13 leads to a global down-regulation of transcription and limits cell cycle progression in glioma cells. We also demonstrate that pharmacological inhibition of CDK12/CDK13 profoundly reduces the *in vitro* and ex *vivo* proliferation of glioma cells and reduces tumor burden in a subcutaneous mouse xenograft model. CDK12/13 inhibition leads to a therapeutically attractive outcome by exploiting a transcriptional addition without directly targeting DNA replication machinery. Identification and further characterization of small molecule inhibitors targeting CDK12/CDK13 with improved pharmacological properties, in particular the ability to cross the BBB, therefore would have a large therapeutic potential for glioblastoma treatment. The potential of CDK12/13 inhibitors for glioblastoma treatment should be investigated further.

## Materials and methods

### Cell culture

All the GSCs used are IDH-wt. GSCs G7, G144, G166, G14, G25, G26 and G30 were a kind gift from S.M. Pollard. GSC cell lines U3013 and U3017 were acquired from HGCC, Uppsala (38). GSC cell lines P3, BG5 and BG7 were a kind gift from Rolf Bjerkvig (39). All GSC cells were cultured in neural stem cell (NSC) medium supplemented with EGF and FGFb as previously described(40). U87MG, Hela, U2OS and HEPG2 cell lines were cultured in DMEM with FBS. MCF10a and breast cancer cell lines, MCF-7, SKBR-3 and MDA-MB-231 were kind gifts from Prof. Ragnhild Eskeland, University of Oslo and Dr. Gunnhild Mælandsmo, Oslo University Hospital, Oslo.

### Cell viability assays

**MTT assay;** 3K-10K cells were seeded in 96-well plates and treated with inhibitors. After 72 hours (h), the MTT assay (Merck, 11465007001) was performed, according to manufacturer’s instructions. **Proliferation assay;** 200,000 cells were seeded in duplicate in 6-well plates. Cells were counted with trypan blue and replated every 3-4 days. **Clonogenic survival assay;** 10,000 cells were seeded in duplicate in 6-well plates. After 14 days, the medium was removed and cells were given a wash with PBS. Crystal violet staining (0.05% Crystal violet, 1% CH_2_O, 1% MeOH in PBS) was added for 20 minutes (min), after which the cells were rinsed with water and left to air dry. ***Ex vivo* assay** was performed on standardized 3D patient-derived organoids from glioblastoma IDH-wt patients that were reformed from isolated single tumor cells as previously described (24, 25). More information in supplemental methods. **Competition assays** were performed as described in supplemental methods.

### Cell migration experiment

Migration patterns of GSCs were studied in 96-well glass bottom plates (Greiner Sensoplate (M4187-16EA, Merck)) after treatment with DMSO control, 500 nM THZ531 or 1.0 µM Gefitinib overnight. More information is provided in the supplemental methods section.

### Immunohistochemistry for CDK12 expression in human glioblastoma tissue

Immunohistochemistry (IHC) for CDK12 expression was performed with polyclonal rabbit-anti-human-CDK12 antibody (ab246887; Abcam; dilution 1:50) using the protocol described in supplementary methods section. To determine the expression of CDK12 in human patients, we assessed glioblastoma and control tissue for immunoreactivity to CDK12 (Figure 2A and Supplementary Figure 2A). A brief description of patient characteristics is included in supplemental Table 1.

### Mouse experiments

SR-4835 (MedChemexpress) was dissolved in 10% DMSO / 90% (30%) Hydroxypropyl-b-Cyclodextrin (hp-BCD) and TMZ (Selleckchem), was dissolved in 10% DMSO in distilled water and administered per os (PO) using gavage. Weekly SR-4835 dosage was five days daily in the week with a two-day break whereas TMZ dosage was daily. For the combination treatment, compounds were dosed at the same time with TMZ dosed first, followed by SR-4835. More information is provided in the supplemental methods section.

### Ethics

For glioma organoids, patient samples were collected from patients having given informed consent, and ethical approval has been obtained from the research ethics committee in Luxembourg (National Committee for Ethics in Research (CNER), as described (24). The use of fully anonymised human tissue (IHC) was also approved by CNER (reference number 1121-278). Mice experiments were carried out by Crownbio, UK and animal welfare were complied with the UK Animals Scientific Procedures Act 1986 (ASPA) in line with Directive 2010/63/EU of the European Parliament and the Council of 22 September 2010 on the protection of animals used for scientific purposes.

#### SR-4835 dose response

3K-5K cells were seeded in 96-well plates and next day, were treated with 3.16-fold dilution in triplicate (31.6µM, 10µM, 3.16µM, 1µM, 316nM, 100nM, 31.6nM, 10nM and 3.16nM). After 72h, CellTiter-Glo assay (Promega) was performed, according to manufacturer’s instructions. The IC50 data used for Figure 3F is available on request.

### Immuno-blotting

Cells were lysed directly in 1.25X Laemmli sample buffer, sonicated and denatured at 95°C for 5 minutes. Samples were loaded and the protein separated in Novex tris-glycine 6% gels (Life Technologies, XP00062BOX) and transferred to nitrocellulose membranes, subsequently standard immuno-blotting procedures were followed with details provided in the supplemental methods section.

### SLAM-seq and CUT&RUN

Cells were treated with THZ531 or vehicle control before they were harvested and were processed using the protocol described in (29),(28), with details described in the supplemental methods section.

### Bioinformatics and data processing

Please see the supplemental methods section for details on SLAM-seq data processing, CUT&RUN and visualization methods.

### Flow cytometry

Cell cycle changes were analysed using the Click-iT EdU Alexa Fluor 647 Flow Cytometry Assay Kit (Invitrogen, CA, USA). Cells were labeled with EdU for 3 h after drug treatment with THZ531. Dead cells were marked using LIVE/DEAD Near-IR (Life Technologies, L10119), and Hoechst was used to mark DNA. Flow cytometry was performed using BD Fortessa (BD Biosciences, CA, USA). Analyses were made using the Flowjo software (FlowJo, LLC, OR, USA).

### Data Availability

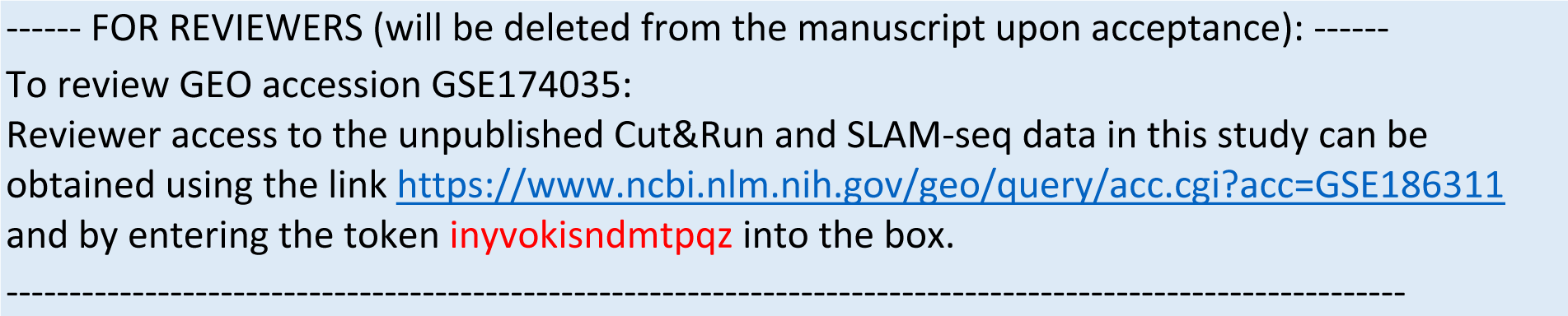

All Cut&Run and SLAM-seq data are deposited at NCBI’s Gene Expression Omnibus (41) under the accession number GSE186311.

## Acknowledgements

We would like to thank the AML and Flow Cytometry Core Facility at Oslo University Hospital for help with microscopy and flow cytometry experiments.

## Declaration

The authors declare no potential conflicts of interest.

## Supplemental methods

### Lentivirus production and transduction

For lentivirus production, 293FT cells were transfected with VSV, Pax8 and expression constructs, using lipofectamine. After 8h, cells were washed and cultured in NSC medium. After 48h, the medium was collected and passed through a 0.45 μm filter. For transduction, GSCs were cultured in NSC medium containing virus particles supplemented with 8 μg/ml polybrene. After 48h of transduction, cells were passaged and cultured in selection medium for 2 days.

### Competition assay

Stably expressing Cas9 G7 cells are derived by transducing GSC G7 cells with lentivirus produced from lentiCas9-Blast (Addgene #52962). The sgRNAs were expressed from U6-sgRNA-SFFV-puro-P2A-EGFP, which is derived from (Addgene: #57827) and reported here ^25^. The sgRNA sequences are available on request. The stably expressing Cas9 G7 cells were transduced with the sgRNAs expressing lentiviruses to achieve 40-60% GFP expressing cells. The transduced cells were subsequently cultured for a period of 21 days and population of sgRNA expressing GFP+ cells were analyzed at indicated days by flow cytometry.

### Cell migration experiment

Migration patterns of human GSCs were studied in 96-well glass bottom plates (Greiner Sensoplate (M4187-16EA, Merck)) after treatment with DMSO control, 500 nM THZ531 or 1.0 µM Gefitinib overnight. Live cell imaging was performed using an ImageXpress Micro Confocal High-Content microscope equipped with an environmental control gasket, maintaining 37°C and 5% CO_2_, and controlled by the MetaXpress 6 software (Molecular Devices). Images were acquired in widefield mode using a 20x 0.45 NA Ph1 air objective, a phase contrast ring, and transmitted light for visualizing the contour of cells. Cell migration was registered for a total time of 8 h (h). For each well 2 sites were imaged with a time interval of 4 minutes (min) between frames. Acquired time lapse movies were analyzed using the TrackMate ^25^ plugin in the Fiji ImageJ software ^26^ and *in-house* Python-based scripts (Python 3.7.6).

***Ex vivo* assay** were performed on standardized 3D patient-derived organoids that were reformed from isolated single tumor cells as previously described {Golebiewska, 2020 #163; Oudin, 2021 #164}. For the CDK12/CDK13 inhibitors, four patient-derived GBM IDHwt tumor cells, exhibiting a typical range of key GBM genetic alterations including deletion of CDKN2A/B, amplification of CDK4/6 and EGFR, mutations IN TP53, PTEN, PIK3CA, EGFR were selected. Briefly, isolated tumor cells were seeded in a 384-well plate at a density of 1000 cells/well and were cultured on an orbital shaker for 72 hours to reform organoids prior to inhibitor treatment. 3D organoids were cultured in a volume of 25 µl DMEM medium/well supplemented with 10% FBS, 2mM L-Glutamine, 0.4mM NEAA, and 100U/ml Pen/Strep (all from Lonza) at 37°C under 5% CO_2_ and atmospheric oxygen. Organoids were treated with the following inhibitors: THZ531 (CDK12/13), SR4835 (CDK12/13), Abemaciclib (CDK4/6) and Lomustine from MedChemExpress in a twofold and six-point serial dilution series ranging from 16 µM to 500 nM (THZ531), 4µM to 125nM (SR4835) and 32 µM to 500 nM (Abemaciclib and Lomustine) from a stock solution of 10 mM prepared in dimethyl sulfoxide (DMSO) (Sigma-Aldrich). A set of control wells with cells treated with DMSO was included on all the plates and the assay was performed with three technical replicates. After 3 days (72 hours) of incubation with the inhibitors, cell viability was measured using the commercially available CellTiter-Glo® 3D Cell Viability Assay (Promega) according to the manufacturer’s instructions. Luminescence was measured with a ClarioStar plate reader (BMG Labtech). The relative cell viability for each inhibitor was calculated by normalization to DMSO control per condition. Dose response curves (DRCs) were fitted by nonlinear regression analysis using GraphPad Prism software: best-fit lines and the resulting IC_50_ values were calculated using log [inhibitor] versus normalized response—variable slope (four parameters).

### Immunohistochemistry for CDK12 expression in GBM patients

Anonymized formalin-fixed and paraffin-embedded (FFPE) tissue samples from routine diagnostics were subjected to CDK12 immunohistochemistry. Minimal clinical data such as age, sex and final pathological diagnoses were included (see **supplementary table-1**). In brief, immunohistochemistry was performed on 2-3 μm thick slices of glioblastoma tissue (n = 5 patients) and on CNS tissue without glioblastoma (n = 2 patients) using an automated IHC staining system Dako Omnis (Agilent, Santa Clara, California, USA). For further details, please see supplementary table-1. Tissue from a patient with fasciitis was used as control tissue. The staining procedure included heat and chemical treatment of the slides with EnVision FLEX TRS at low pH at 97°C (20 min), incubation (30 min) with polyclonal rabbit-anti-human-CDK12 antibody (ab246887; Abcam; 1:50) and 3 min endogenous enzyme block with EnV FLEX Peroxidase-Blocking solution. Signal enhancement was achieved by incubating slides with EnV Flex + Rabbit LINKER (10 min). EnV FLEX/HRP labeled polymer (20 min) was used as secondary antibody, followed by incubation with the EnVision FLEX DAB Chromogen for 5 minutes. Slides were counterstained with hematoxylin and mounted with coverslipping film Tissue-Tek (Sakura, Staufen, Germany). Slides were then scanned in the Hamamatsu NanoZoomer 2.0-HT (Hamamatsu Photonics), digitized, and transferred to a computer screen. Brightness, gain, and contrast were all kept constant during image acquisition. Glioblastoma and CNS tissue as well as control tissue were then assessed for immunoreactivity (negative, weak, moderate or strong) to CDK12 (**see also Figure 2A/Supplementary figure 2A**).

**Supplementary Table 1 :** Patients characteristics of the samples included

| Patient No. | Slide No. | Age (y) | Gender | Tissue | Diagnosis |
| --- | --- | --- | --- | --- | --- |
| 1 | C12 | 81 | female | Fascia | fasciitis |
| 1 | C13 | 81 | female | Fascia | fasciitis |
| 2 | C18 | 40 | male | CNS | Cavernous hemangioma |
| 3 | C14 | 59 | female | CNS | Glioblastoma, CNS WHO grade 4 |
| 4 | C15 | 74 | female | CNS | cerebral amyloid angiopathy (CAA) |
| 5 | C16 | 85 | female | CNS | glioblastoma, CNS WHO grade 4 |
| 6 | C17 | 69 | female | CNS | glioblastoma, CNS WHO grade 4 |
| 7 | C19 | 64 | male | CNS | glioblastoma, CNS WHO grade 4 |
| 8 | C21 | 55 | male | CNS | rest/recurrent glioblastoma, CNS WHO grade 4 |

### Mouse experiments

SR-4835, from MedChemexpress was dissolved in 10% DMSO / 90% (30%) Hydroxypropyl-b-Cyclodextrin (hp-BCD) and TMZ from Selleckchem, was dissolved in 10% DMSO in distilled water and administered per os (PO). Weekly SR-4835 dosage was five days daily in the week with a two days break whereas TMZ dosage was daily. For the combination treatment, compounds were dosed at the same time with TMZ dosed first, followed by SR-4835. For a more detailed A subcutaneous U87-MG-luc model was established by Crown Bioscience Inc. (Leicester, UK) using the parental U87-MG cell line from ECACC, transduced in house to express luciferase. Animals were housed in IVC housing with a 12 h light/dark cycle and access to Teklad 2919 and sterile water ad libitum. **Tolerability experiment** Athymic nude mice aged 7-8 weeks old from Envigo were dosed with 20 mg/kg and 30 mg/kg SR-4835 for 2 weeks with the weekly cycle five days on and two days off. **Efficacy experiments** The *in vivo* efficacy of the SR-4835, either as a monotherapy or in combination with Temozolamide, was further evaluated in the clinically relevant subcutaneous CDX U87-MG-luc xenograft model. These experiments were conducted at Crown Bioscience, Inc. (Leicester, UK) in 7-8 week old athymic nude mice (Envigo, UK). Forty-eight mice were enrolled in the efficacy study, eight mice per cohort for six cohorts. Eight million U-87-MG-luc cells were injected subcutaneously injected into athymic nude mice acquired from Envigo. All animals were randomly allocated to the six different study. Randomization was performed on day nine post injection, prior to the treatment start. The treatments were undertaken for 2 weeks with the five day on, 2 day off cycle per week for SR-4835 and daily for TMZ. Tumors were measured 3 times a week and tumor volumes were estimated by measuring the tumor in two dimensions using electronic callipers. **Humane Endpoints** Any mouse with tumor volume/measurement at terminal size (e.g. mean diameter 15mm) was terminated. After one measurement of body weight loss (BWL) > 10% on the day of dosing, a treatment break was given and treatment resumed when the body weight recovered to BWL < 5% (compared to day 1 treatment). Any mouse with BWL > 15% for 3 consecutive measurements (compared to day 1 treatment) was euthanized and any mouse with body weight loss ≥ 20% was Schedule 1 culled.

### Immuno-blotting

Cells were lysed directly in 1.25X Laemmli sample buffer, sonicated and denatured at 95°C for 5 minutes. Samples were loaded and the protein separated in Novex tris-glycine 6% gels (Life Technologies, XP00062BOX) and transferred to nitrocellulose membranes. Membranes were incubated in blocking buffer (5% FBS in TBST (TBS with 0.2% Tween-20) for 1.5h at room temperature and incubated with primary antibody (dilutions are indicated in the table below) in the blocking buffer overnight at 4°C. Membranes were given three washes in TBST for 10 minutes, then incubated 35 minutes with appropriate secondary antibodies in blocking buffer and washed for another 3×10 minutes. Chemiluminescent detection was performed using Chemiluminescent substrate kit from Fisher Scientific, catalog 34580.

**Supplementary Table 2 :** Antibody dilution

| Antibody | Manufacturer | Catalog number | Dilution for CUT&RUN | Dilution for WB |
| --- | --- | --- | --- | --- |
| RNAPII total | Mbl | MABI0601 | 1:100 | 1:10 000 |
| RNAPII pSer2 | Mbl | MABI0602 | 1:100 | 1:10 000 |
| RNAPII pSer5 | Mbl | MABI0603 | 1:100 | 1:10 000 |
| H3K27ac | CST | D5E4 | 1:100 | 1:1000 |
| H3K27me3 | CST | C36B11 | 1:100 | 1:1000 |
| RNAPII pThr4 | Active Motif | 61461 | NA | 1:1000 |
| RNAPII pSer7 | Active Motif | 61087 | NA | 1:1000 |
| Vinculin | Sigma | V9131 | NA | 1:10 000 |
| PARP | CST | 9542 | NA | 1:1000 |
| GAPDH | Santa Cruz<br>Biotechnology | sc25778 | NA | 1:20 000 |
| IgG | abcam | ab6721 | 1:100 | NA |

**qPCR primers** are available on request

### SLAM-seq

SLAM-seq was performed according to the previous described protocol ^26^. Briefly, cells were seeded at approximately 70% confluency, and treated with THZ531 or vehicle control. 1 hour before harvest, 4sU was added to the medium at a final concentration of 500 μM. The samples were kept in the dark as RNA was extracted with Qiagen’s RNeasy Plus Mini Kit. 3 μg RNA was alkylated with iodoacetamide (Sigma, 10 mM) for 15 minutes, and the RNA was repurified by ethanol precipitation. 250 ng RNA was used to make libraries with Lexogen’s QuantSeq 3′ mRNA-Seq Library Prep Kit FWD for Illumina and PCR Add-on Kit for Illumina. Deep sequencing was performed using the NovaSeq platform (Illumina).

### CUT&RUN

Cells were treated with THZ531 or vehicle control before they were harvested, washed and bound to Concanavalin A-coated magnetic beads. The cells were then permeabilized with wash buffer (20 mM HEPES, pH7.5, 150 mM NaCl, 0.5 mM spermidine and a Roche complete tablet per 50 ml) containing 0.02% Digitonin. The cell-bead suspension was incubated with 0.5-1 μg of respective antibody in a total volume of 50 μL overnight at 4°C on a nutator. After 3 washes with 1 mL Digitonin buffer, cells were resuspended in 50 μL volume with pAG-MNase and nutated for 10 minutes at RT. Cells were given two washes with Digitonin buffer, chilled on ice, and ice-cold CaCl_2_ was added, before nutating for 2 hours at 4°C. STOP buffer was added (340 mM NaCl, 20 mM EDTA, 4 mM EGTA, 0.02% Digitonin, 50 μg/mL RNAse A, 50 μg/ml glycogen), and tubes were incubated at 37°C for 10 minutes in a ThermoMixer, to release fragments into solution. After centrifugation at 16,000 x g for 5 minutes at 4°C, tubes were placed on a magnet stand and the liquid transferred to new tubes. DNA was extracted using Qiagen’s MinElute PCR Purification Kit, according to manufacturer’s instructions. Quantification was done by Qubit analysis, and libraries were made with 2 ng DNA using the NEBNext® Ultra™ II DNA Library Prep Kit for Illumina” (New England Biolabs, E7645), according to manufacturer’s instructions. Deep sequencing was performed using the NovaSeq platform (Illumina).

### SLAM-seq data processing

3’ UTR annotations were obtained from ^29^. All further processing was done on the Galaxy server, using the SlamDunk pipeline (http://github.com/t-neumann/slamdunk). Prior to mapping, the quality of the sequencing of the reads was inspected using FastQC (v.0.72+galaxy1) (https://www.bioinformatics.babraham.ac.uk/projects/fastqc/). Adapters were trimmed from raw reads using cutadapt through the trim_galore (v.0.4.3.2) wrapper tool with adapter overlaps set to 3bp. The reads were then processed using SlamDunk (v.0.4.1+galaxy2). Settings were adjusted to alignment against the human genome (GRCh38), 12bp trimming from the 5’ end, with multi-mapper retention strategy for 100 alignments, filtering for variants with a 0.2 variant fraction, filtering for base-quality cutoff of ≥27, and filtering for ≥1 T>C conversions.

### Cut&Run data processing

Prior to mapping, the quality of sequenced reads were inspected using FastQC (v. 0.10.1) (https://www.bioinformatics.babraham.ac.uk/projects/fastqc/), fastqScreen (v. 0.11.4, ^41^), and MultiQC (v. 1.7, ^42^) and reads were mapped to hg38 using Bowtie2 (v. 2.2.9, ^43^) and the settings “--local --very-sensitive-local --no-unal --no-mixed --no-discordant --phred33 -I 10 -X 700”, and filtered for an insert size between 20 and 120 base pairs using samtools (v. 1.10, ^44^) in accordance with instructions from the CUT&RUN protocol ^45^ in the following pipe: “samtools view -h -f 66 | awk -F’\t’ ‘function abs(x){return ((x < 0.0)-x : x)} {if ((abs($9) >= 20 && abs($9) <= 120) || $1 ∼ /^@/) print $0}’ | samtools view -Sb – “. Filtered mapped reads were deduplicated and imported into EaSeq v. 1.2 ^46^ using default settings and unless specified subsequent analysis and visualisation was performed using the integrated tools in EaSeq.

### Genome-wide data sources

All Cut&Run and SLAM-seq data are deposited at NCBI’s Gene Expression Omnibus ^47^ under the accession number GSE186311. Refseq gene annotations ^48^ were acquired from the UCSC table browser ^49^. CDK and cyclin genes were identified based on matching the strings ‘CDK’ or ‘CCN’ to gene symbols. RPL/RPS gene symbols were obtained from http://ribosome.med.miyazaki-u.ac.jp/rpg.cgi?mode=orglist&org=Homo%20sapiens, Histone gene symbols were obtained from http://www.informatics.jax.org/mgihome/nomen/gene_name_initiative.shtml, house-keeping gene symbols were obtained from ^50^, DDR-gene symbols were obtained from https://www.mdanderson.org/documents/Labs/Wood-Laboratory/human-dna-repair-genes.html, BRCA-ness gene symbols were obtained from ^51^ and gene symbols for glioma transcription factors and targets were obtained from ^11^.

### Cut&Run and SLAM-seq visualization and integration

Graphs of average Cut&Run signal as well as heatmaps were generated in EaSeq using the ‘Average’ and ‘HeatMap’-tools, respectively. K-means clustering was performed using the ‘Cluster’-tool with the clustering methodology set to k-means, the offset set to +/-1kbp and log-transformation disabled. Output from the SLAM-seq processing was analysed for differential expression using DeSeq2 ^52^ with default settings, and size factors estimated on the total mRNA reads for global normalization. Transcripts were subgrouped according to adjusted p-values from the differential expression analysis, with the group ‘Most down’ having adjusted p-values below 10^−5^ and the group ‘Less down’ having adjusted p-values between 10^−5^ and 0.05. Volcano plots were generated based on log2 fold differences and adjusted p-values from DeSeq2 and visualized using EaSeq and the ‘Scatter’-tool or Microsoft Excel 2016. The number of selected gene subsets found within the significantly regulated genes was counted and compared to that expected by chance using Chi-square testing, and p-values for all shown comparisons were Bonferroni-adjusted before being plotted in bubble diagrams. Cut&Run values for 1D heatmaps of signal at TSSes were quantified using the ‘Quantify’-tool and default settings except for using offsets of+/-1kbp and visualized together with ‘basemean’ and log2 fold difference values from the DeSeq2 output of Nascent transcripts using the ‘ParMap’-tool. The order of the TSSes was determined based on first the grouping as mentioned above, and then the average expression value in all conditions (Basemean). GO-term enrichment analysis of significantly regulated transcripts was done using g:Profiler (https://biit.cs.ut.ee/gprofiler/,53). Bee-swarm plots were made using the using R (https://www.R-project.org/) and the beeswarm package (The Bee Swarm Plot, an Alternative to Stripchart, version 0.2.0, A Eklund (2016), CRAN). Integration of expression data with cell-cycle related transcriptional changes was done using published results ^30^. The ‘polar coordinates’ from transcripts that were found to be significant in their work was used as the X-axis when visualizing the moving average log 2 fold difference in the expression of 100 transcripts (Y-axis).

## Supplementary legends

**Supplementary Figure 1:**
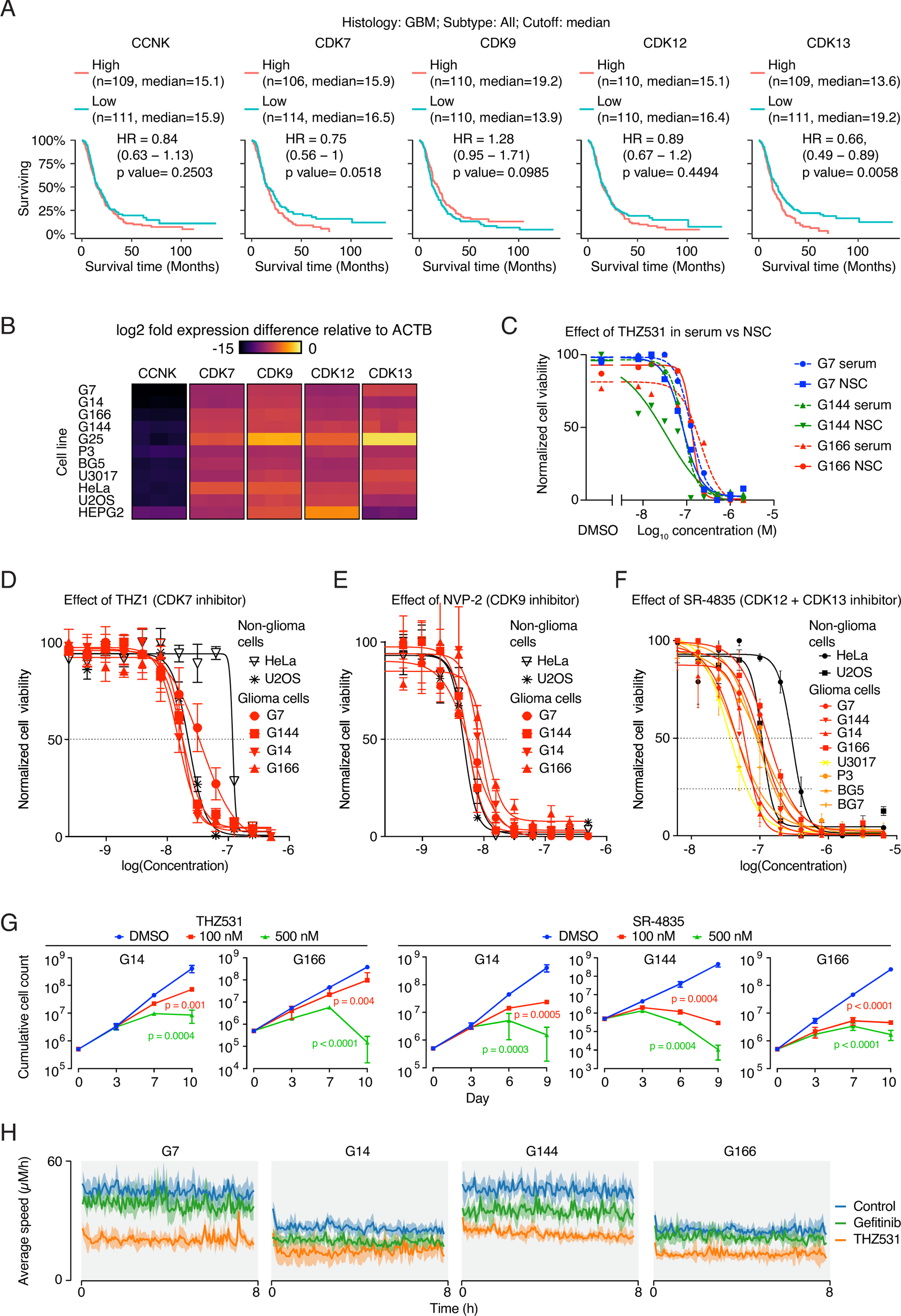
**(A)** Survival analyses from Gliovis showing relation between survival of patients and expression of tCDK used in the study. (**B**) Heatmap showing the mRNA expression of tCDKs in the cell lines used in the study. (C) Three high-grade glioma cell lines were cultured in serum-free or serum-containing media and were treated with increasing doses of THZ531. After 72h, the cell viability was assessed using Cell-Titer-Glo. Graph displays a dose-response curve with percent cell viability relative to the DMSO control for each cell line. Data represent mean ± SD of three replicates. (**D**) Four high-grade glioma cell lines and two non-glioma cell lines were treated with increasing doses of THZ1. After 72h, cells were subjected to the MTT assay. Graph displays a dose-response curve with percent cell viability relative to the DMSO control for each cell line. Data represent mean ± SD of three replicates. (**E**) Four high-grade glioma cell lines and two non-glioma cell lines were treated with increasing doses of NVP-2. After 72h, cells were subjected to the MTT assay. Graph displays a dose-response curve with percent cell viability relative to the DMSO control for each cell line. Data represent mean ± SD of three replicates. (**F**) Eight high-grade glioma cell lines and two non-glioma cell lines were treated with increasing doses of SR-4835. After 72h, cells were subjected to the MTT assay. Graph displays a dose-response curve with percent cell viability relative to the DMSO control for each cell line. Data represent mean ± SD of three replicates. (**G**) *in vitro* cell proliferation assay of GSCs treated as indicated. Data represent mean ± SD of two replicates. (**H**) Effect of THZ531 treatment on the migration of glioma cells. Average speed of migration is plotted over time. Gefitinib, an EGFR inhibitor is used a positive control.

**Supplementary Figure 2:**
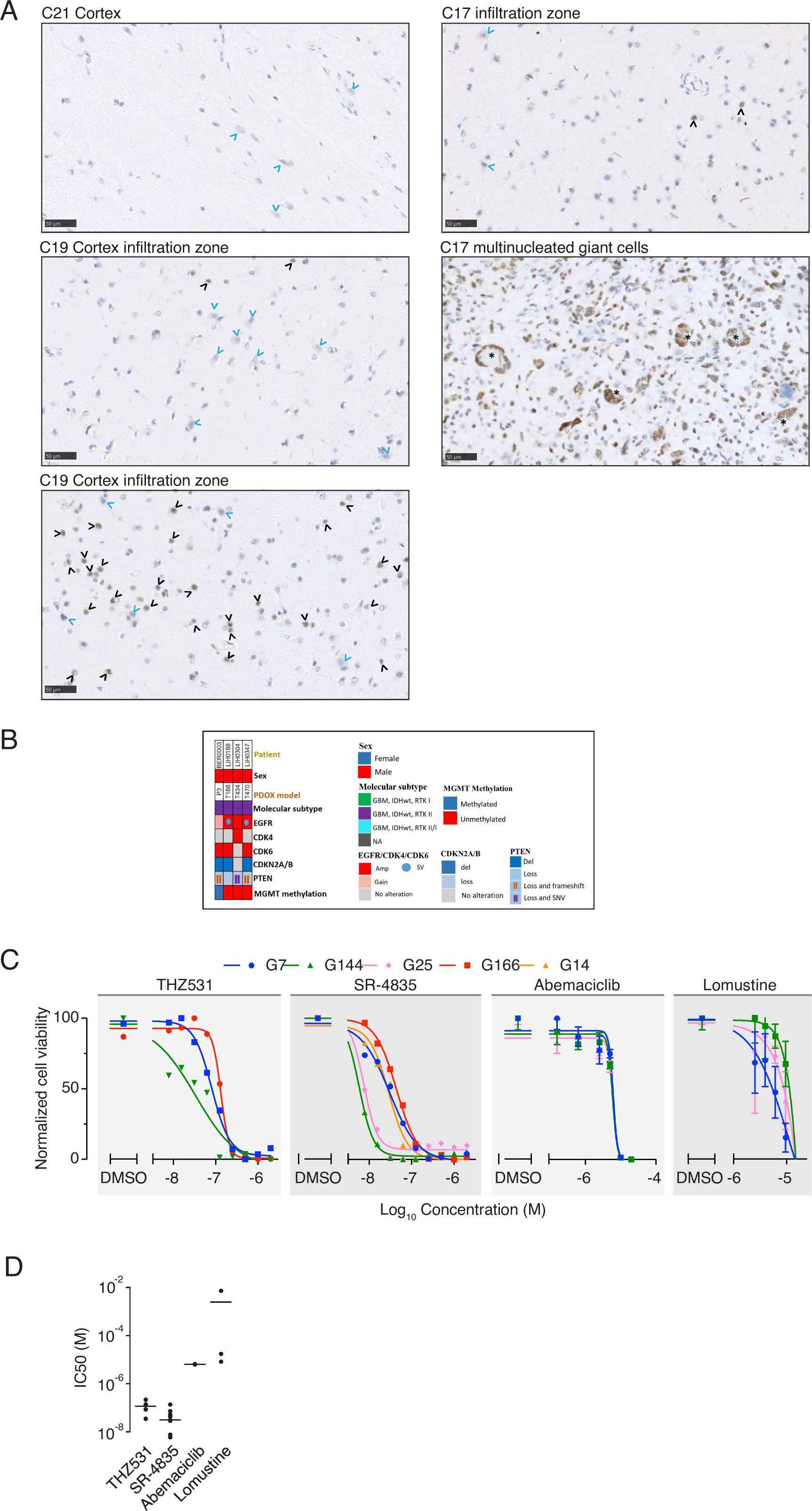
**(A)** Representative images of CDK12 immunohistochemistry in cortex/infiltration zone/cell-rich tumor of glioblastoma patients. Nuclear CDK12 expression absent in cortex areas without obvious tumor cell infiltration (top) while the number of CDK12-positive cells increases with tumor cell density in the infiltration zone (CDK12-positive cells = black arrowhead; CDK12-negative cortical neurons = blue arrowhead). Multinucleated giant cells (asterisk, bottom) in highly cellular areas expressing CDK12. Scale bar 50 µm. **(B)** Summary of the patient characteristics of the ex vivo GBM organoids. **(C)** Four high-grade glioma cell lines were treated with increasing doses of inhibitors as indicated. After 72h, the cell viability was assessed using Cell-Titer-Glo. Graph displays a dose-response curve with percent cell viability relative to the DMSO control for each cell line. Data represent mean ± SD of three replicates. **(D)** Dot-plot showing IC50 values for dose response of inhibitors on a panel of GSCs shown in (C).

**Supplementary Figure 3:**
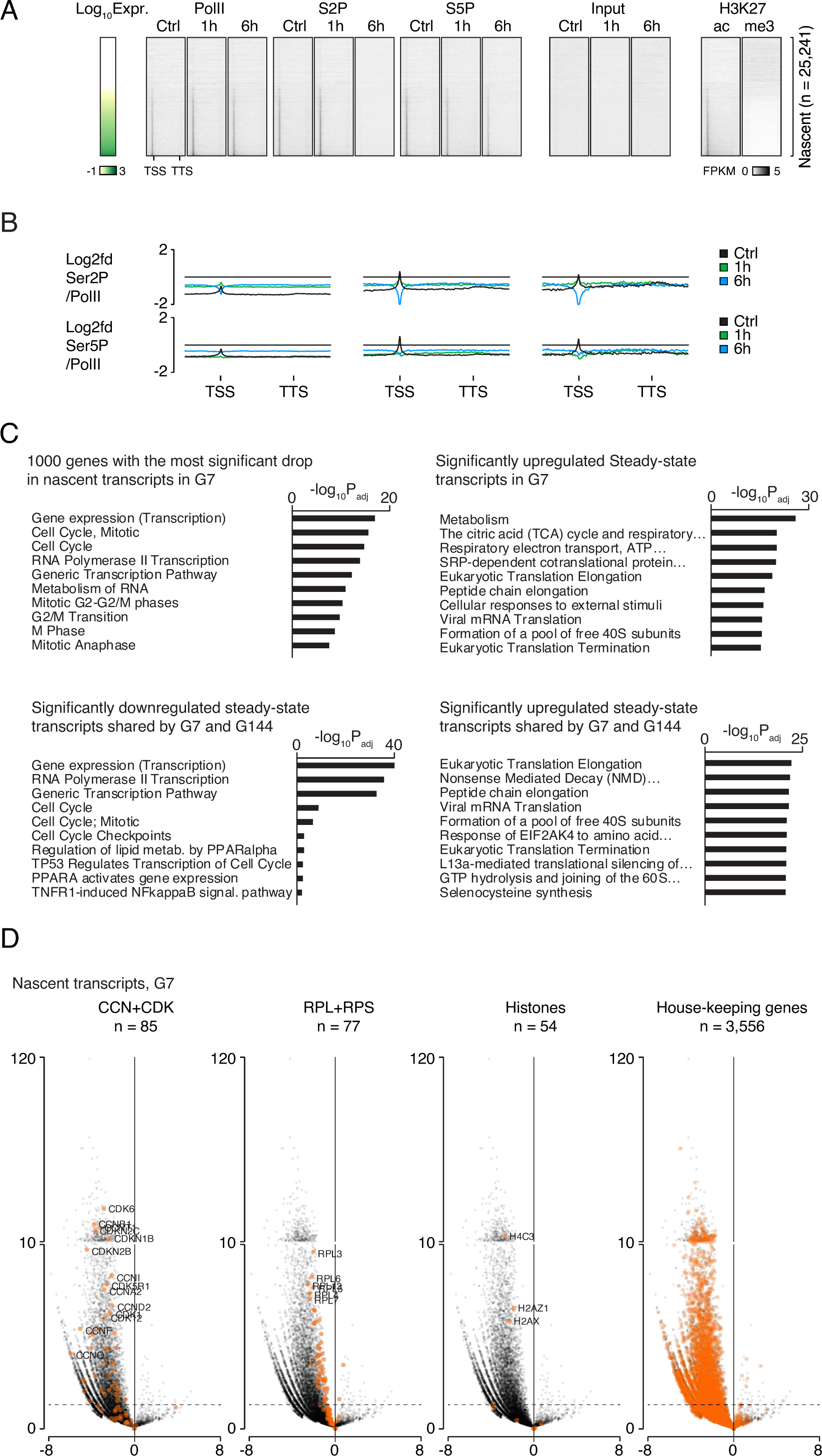
(**A**) Heatmaps of Cut&Run signal from RNAPII, RNAPII phosphorylation states, selected histone mark modifications, as well as input at gene bodies and immediate upstream and downstream regions (+/-25% of gene length) from G7 cells treated with either THZ531 for 1 hr, or THZ531 for 6 hrs as well as DMSO controls. Genes were ordered vertically based on their total expression level. The horizontal extent of each gene and the upstream and downstream regions corresponding to quarter of the gene length is fitted within the same visual space in the heatmaps regardless of its absolute extent. TSS and TTS illustrate transcription start sites and termination sites, respectively. Cut&Run and input levels are FPKM normalized. (**B**) Graphs of average Cut&Run signal from RNAPII phosphorylation states normalized to RNAPII levels at all gene bodies and surrounding loci. RNAPII and RNAPII modification states were obtained as described in Figure 4A. The horizontal extent of each gene and the upstream and downstream regions corresponding to half a gene length is fitted within the same visual space in the heatmaps regardless of its absolute extent. TSS and TTS illustrate transcription start sites and termination sites, respectively. Cut&Run levels are FPKM normalized. (**C**) Bar diagrams showing the most significantly enriched gene ontology (GO) terms from selected subsets of genes from steady-state and nascent RNA. X-axes represent −log10 p-values adjusted for multiple testing. (**D**) Volcano plots showing the overall transcriptional differences in nascent transcripts as in Figure 3E, but with certain gene populations highlighted. X-axes shows the log2 fold difference in transcription in G7 cells treated with THZ531 for six hours compared to DMSO controls. Y-axes shows the −log10 transformed p-values Benjamini-Hochberg corrected for multiple testing. Coloured dots illustrate the transcriptional changes of the listed gene populations.

**Supplementary Figure 4:**
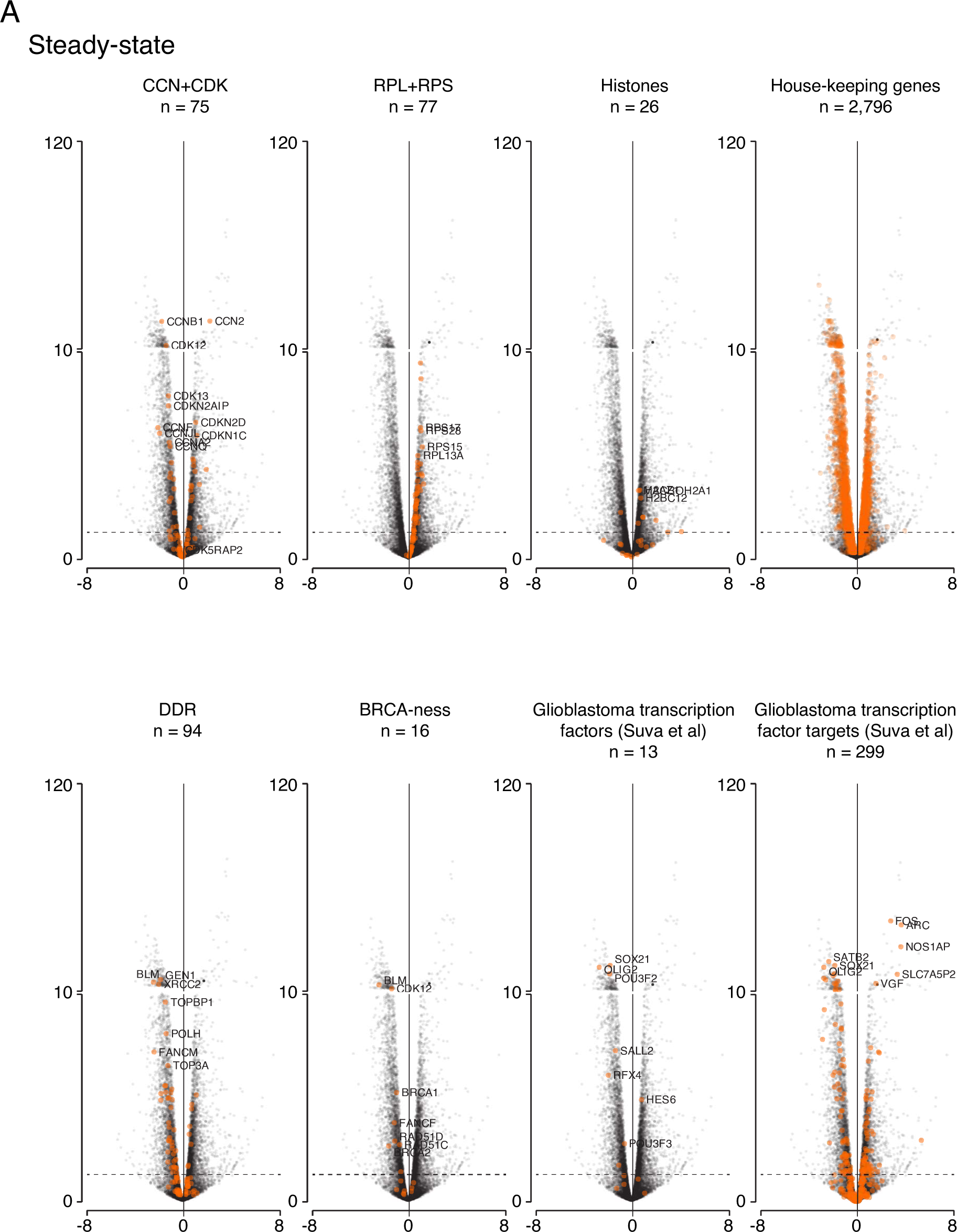
(**A**) Volcano plots showing the overall transcriptional differences in steady-state transcripts as in Figure 3D, but with certain gene populations highlighted. X-axes shows the log2 fold difference in transcription in G7 cells treated with THZ531 for six hours compared to DMSO controls. Y-axes shows the −log10 transformed p-values Benjamini-Hochberg corrected for multiple testing. Coloured dots illustrate the transcriptional changes of the listed gene populations.

**Supplementary Figure 5:**
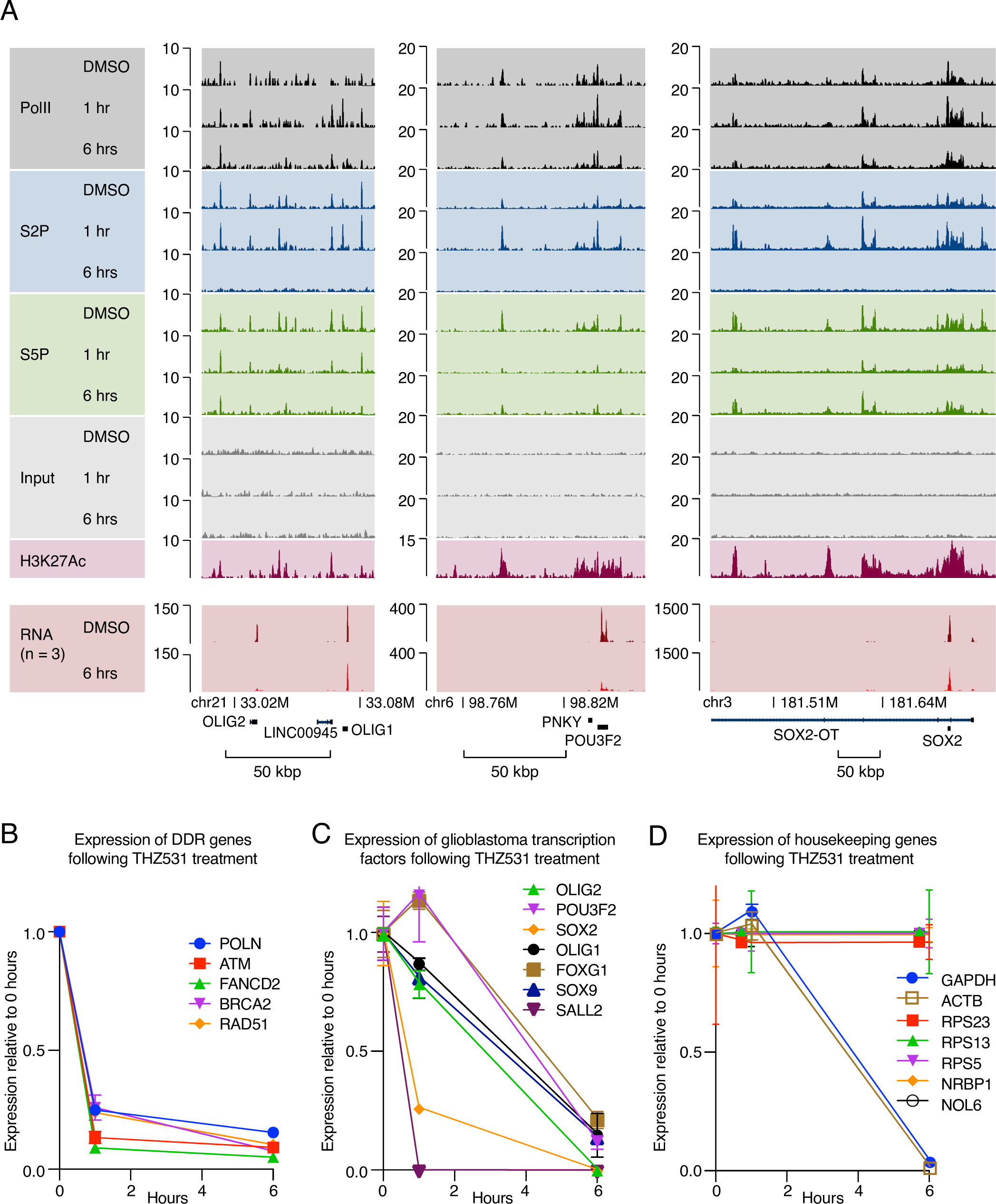
(**A**) Genome browser tracks of Cut&Run signal density and total transcript-levels (bottom) at the Olig2, Pou3f2 and Sox2 loci at indicated treatments. (**B-D**) RT-qPCR analyses of the mRNA levels of DDR genes (**B**), glioblastoma transcription factors (**C**) or housekeeping genes (**D**) in G7 cells treated with 500 nM THZ531 for 1 h and 6 h. Data represent mean ± SD of two replicates.

**Supplementary Figure 6:**
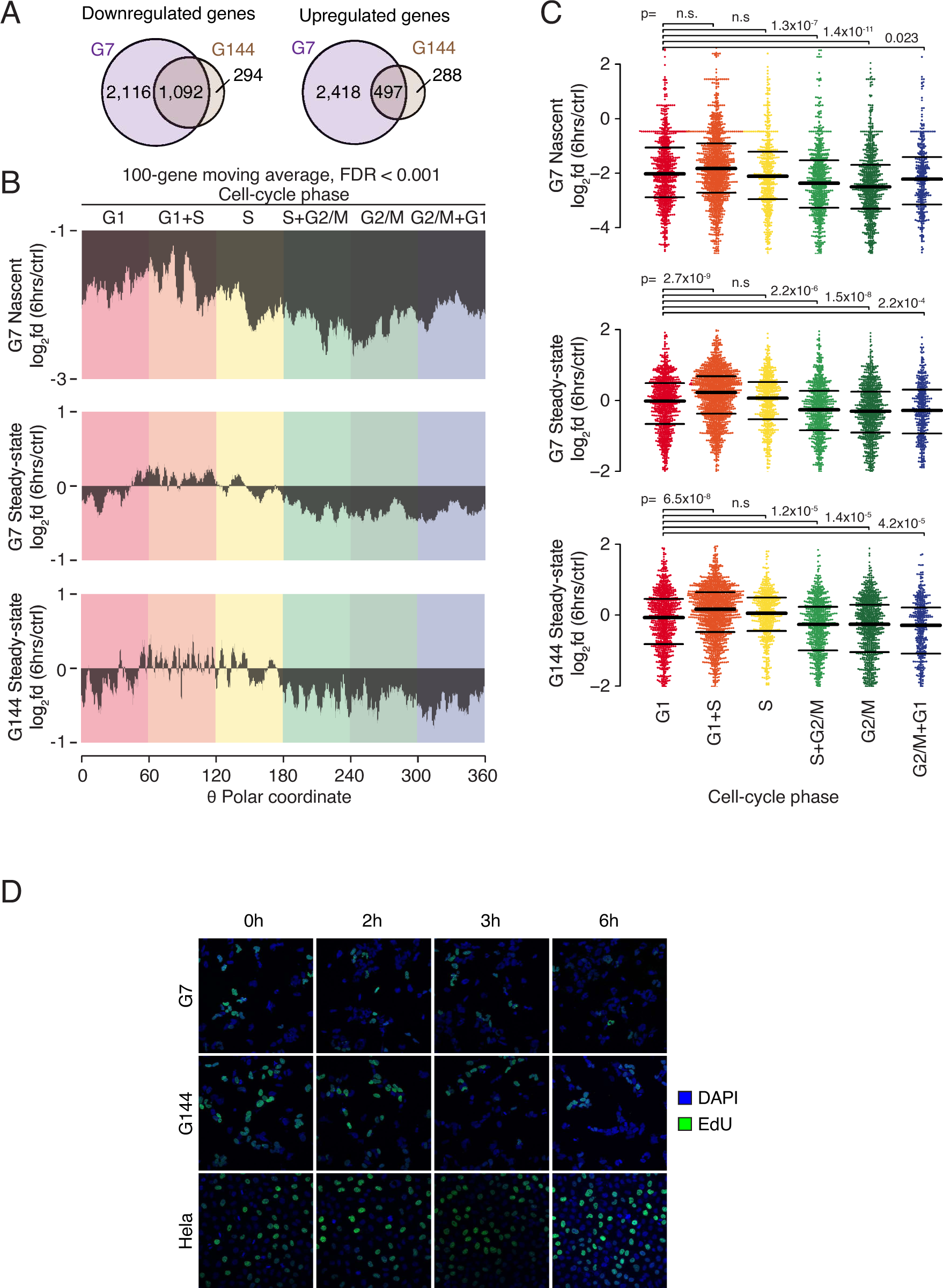
(**A**) Venn-diagrams illustrating the overlap in the populations of genes being downregulated (left) as well as upregulated (right) in G7 cells compared to G144 cells. (**B**) Graphs illustrated the moving average of transcriptional changes in nascent (top) or steady-state (middle, bottom) transcripts from G7 (top, middle) or G144 (bottom) cells treated with THZ531 for 6 hours compared to DMSO controls. Transcripts were ordered according to previously published classification of cell-cycle timing. Only transcripts, for which transcriptional timing could be assessed with an FDR-value of 0.001 or better, were included. (**C**) Beeswarm-plots of transcriptional changes as in **(B)** with transcripts grouped into six overall groups based on previously published transcriptional timing. P-values were obtained using Mann-Whitney U-tests and Bonferroni-corrected for multiple testing. (**D**) Cells were treated with 500 nM THZ531 at indicated times and EdU incorporation was performed in the last 1 hour by adding 10 µM EdU. Cell staining was done using Click-IT chemistry according to manufacturer’s instructions.

**Supplementary movie 1 and 2:** The movies include raw data with TrackMate overlay to show particle tracking of migrating G7 **(Supplementary movie 1)** and G144 cells (**Supplementary movie 2)**. Prior to imaging, 6000 cells were seeded in wells of a 96-well glass plate and cells exposed to either DMSO (left panel) or 500 nM THZ531 (right panel) overnight. Image acquisition was carried out using a 20x air objective, a time interval of 4 min and a total imaging period of 8 h. The movie has been reduced to show 50 frames. Scale bar is 10 µm.

